# A Single-Aliquot, Enrichment-Free Workflow for High-Throughput Plasma Proteome and N-Glycoproteome Profiling

**DOI:** 10.1101/2025.11.11.687106

**Authors:** Kun-Hao Chang, Jared Deyarmin, Tiara Pradita, Yi-Ju Chen, Stephanie Samra, Gee-Chen Chang, Chong-Jen Yu, Yi-Shuang Chuang, Tabiwang Arrey, Hsiang-En Hsu, Yue Xuan, Pei-Rong Huang, Kuen-Tyng Lin, Ke-Hsin Yen, Irene-Ya Tai, Hsing-jui Tsai, Daniel Hermanson, YaHsuan Chang, Sung-Liang Yu, Pan-Chyr Yang, Yu-Ju Chen

## Abstract

High cancer mortality rates highlight an urgent need for early detection. Plasma proteomics and glycoproteomics provide minimally invasive routes for biomarker discovery, yet achieving an optimal balance among profiling depth, throughput, and longitudinal reproducibility remains a major challenge for clinical application and translation. To overcome this bottleneck, we present a single-aliquot, enrichment-free, paired-run dual-omics pipeline that concurrently profiles the global plasma proteome and N-glycoproteome from unenriched plasma. Following depletion of top14 abundant plasma proteins, we implemented sequential 23-min narrow-window data-independent acquisition (DIA) and 42-min stepped-collision-energy data-dependent (SCE-DDA) runs from the same plasma digest, delivering a clinical throughput of ∼24 patients/day with deep proteome coverage of 3,756±413 protein groups (PGs) and 1,226±78 glycopeptides per sample, including 303 FDA-approved drug targets. Cross-platform benchmarking with a previous generation instrument demonstrated significantly faster (>10-20 fold) profiling speed to achieve 113 PGs/min and high protein abundance reproducibility (Pearson r > 0.9), confirming cross-instrument transferability. Application to a 300-participant lung cohort (cancer, LDCT-detected non-cancer nodules, and controls) revealed differential expression of S100 and annexin family proteins between cancer and nodules. Paired glycoproteomic analysis (n=30) identified site-specific N-glycosylation alterations in FN1, IGHG2, C3, and MET independent of total protein abundance, uncovering additional biomarker candidates for early lung cancer detection. Together, this dual-omics strategy enables deep, scalable, and reproducible plasma analysis, supporting longitudinal biomarker discovery and validation across instruments and laboratories.

## INTRODUCTION

Cancer biomarker discovery presents one of the urgent needs in clinical proteomics due to insufficient detection at early stages and high mortality rates. Nearly all FDA-approved cancer biomarkers are proteins, especially secreted glycoproteins. Thus, the plasma proteome and especially the plasma glycoproteome hold considerable significance for biomarker research. Plasma contains thousands of proteins spanning more than ten orders of magnitude in concentration, with glycoproteins forming a major functional subset involved in signaling, immunity, and metabolism [1,2]. Comprehensive profiling of plasma proteins and their site-specific glycosylation degree can provide insight into disease mechanism, progression, and response to therapy [3,4,5,6].

Despite the wide clinical applications of the plasma proteome and glycoproteome, the field is facing several challenges. Striking the balance between the profiling speed and the proteome depth remains a central issue in plasma biomarker discovery. Classical high-abundance proteins such as albumin and immunoglobulins account for over 90% of total plasma protein mass, while more than thousand proteins collectively contribute less than 10% [7]. This extreme dynamic range produces a strong masking effect, hindering the detection of low-abundance but biologically informative proteins. These trace proteins, such as cytokines, interleukins, or tissue leakage proteins, represent promising biomarker candidates for disease detection or patient stratification [8,9]. Therefore, a robust plasma proteomic workflow capable of achieving a deep proteome landscape is essential for translational research. For clinical implementation, additional challenges include ensuring rapid turnaround time and regulatory-grade reproducibility across time, instruments, sites, and cohorts. These demands are further compounded by multi-omic profiling, which often requires independent workflows for global proteomics and post-translational modification (PTM)-focused analyses.

Over the past decade, methodological innovations have substantially advanced the sensitivity, depth, and throughput of plasma proteomics. One major strategy to enhance detection of low-abundance proteins is high-abundant protein (HAP) depletion, which reduces ion suppression during mass spectrometry (MS) analysis by removing dominant plasma components such as albumin and immunoglobulins through the use of immunoaffinity-based columns or spin filters [10]. When coupled with modern MS instruments, such depletion workflows can enable identification of over 1,500 proteins spanning eight orders of magnitude in dynamic range from microliter-scale plasma input [11]. More recently, nanoparticle-based enrichment has emerged as an alternative approach. Functionalized nanoparticles interact with plasma proteins through electrostatic and affinity forces to form a protein corona, capturing low-abundance proteins without prior depletion. This strategy can identify ∼5,000 proteins from particle-based enrichment processed plasma [12,13]. The depletion strategy required relatively lower plasma volume and cost without predefining the analysis panel [14]. Nevertheless, these approaches highlight the conceptually complementary sample processing method to expand proteome depth with scalability for translational studies.

Plasma glycoproteomic analysis traditionally requires complex enrichment procedures and specialized MS acquisition to address challenges posed by the broad dynamic range of plasma proteins, structural glycan heterogeneity, and low glycopeptide detectability. Conventional workflows typically involve protein depletion and glycopeptide enrichment to achieve sufficient coverage. Yong *et al*. introduced Glyco-CPLL, integrating combinatorial peptide ligand libraries (CPLL), high-pH reversed-phase prefractionation, and HILIC enrichment, identifying 1,644 N-glycopeptides (369 glycoproteins) from 100 µg plasma peptides across three fractions [15]. Yong et al. introduced Glyco-CPLL, integrating combinatorial peptide ligand libraries (CPLL), high-pH reversed-phase pre-fractionation and HILIC enrichment, identifying 1644 N-glycopeptides (369 glycoproteins) from 100 ug plasma peptides across three fractions [15]. Shu *et al*. further advanced coverage with the pMAtchGlyco strategy, reporting 22,677 N-glycopeptides (526 glycoproteins) from 400 µg serum peptides across ten fractions, showing the most comprehensive serum N-glycoproteome to date [16]. However, such depth depends on large sample input and extensive fractionation, often impractical for clinical studies [16]. Most recently, Jager et al. introduced the nGlycoDIA, a narrow-window DIA acquisition on the Thermo Scientific™ Orbitrap™ Astral™ mass spectrometer (MS), achieving ≥3,000 unique glycopeptides (181 glycoproteins) across combined four replicates using 10 µL enriched plasma in a 40-min run (NCE 35) [17]. When analyzed without enrichment under identical conditions, detection dropped to ∼400 glycopeptides across four replicates [17]. In contrast, oxonium ion-based scanning DIA approach (OxoScan-MS) bypasses the depletion and enrichment steps, yielding a total of 1102 glyco--precursors from 5 uL plasma of COVID-19 patients (n=30) and healthy controls (n=15). [18]. Together, these advances illustrate an ongoing trade-off between analytical depth and workflow scalability, as enrichment-based methods continue to provide the greatest depth, while enrichment-free approaches are promising routes for streamlined, clinically adaptable workflow.

Parallel characterization of plasma proteome and glycoproteome provides complementary insights: changes in protein abundance often highlight disease-related pathways, while glycosylation alterations provide higher specificity for disease subtypes and may occur independently of total protein levels. Thus, integrated proteomics and glycoproteomics likely expands the biomarker discovery space with higher sensitivity in differentiating disease phenotypes. To offer both layers of information, we introduce a single-aliquot, enrichment-free, paired-run DIA pipeline for concurrent plasma proteome and N-glycoproteome profiling. We adopted a top14 protein depletion-based strategy that requires only microliter-scale plasma volumes and avoids the capture bias associated with nanoparticle enrichment, thereby preserving a more comprehensive and unbiased plasma proteome for dual-omics analysis. Leveraging the high acquisition speed of the Thermo Scientific™ Orbitrap™ Astral™ Zoom mass analyzer, we implemented sequential 23-min DIA and 42-min SCE-HCD-DDA runs from the same sample digest. This workflow achieves a balance between proteome depth and speed, demonstrates good reproducibility (median CV <10%), shows high cross-platform correlation with data from the same set of samples generated on a Thermo Scientific™ Orbitrap™ Fusion Lumos MS (r >0.90), and supports clinical-scale profiling (∼24 patients/day). We applied this approach for biomarker discovery in a low-dose CT (LDCT) screening cohort.

## METHODS

### Plasma collection

Plasma samples were allotted from the de-identified and archived Formosa and Trio low-dose computed tomography (LDCT) trial participants, including normal (n=125), benign tumor patients (n=125), and malignant tumor patients (n=50). This study was conducted under Institutional Review Board (IRB) approval at National Taiwan University Hospital (IRB number: 201410013RIND) and Chung Shan Medical University Hospital (IRB number: CS1-21190). Pooled plasma samples were prepared from the 300 plasma samples for analytical and longitudinal quality control assessment. All the samples were stored in a −80°C freezer before sample preparation.

### Sample preparation

For depletion of abundant serum proteins, 200 µL of High-Select∼ Top14 Abundant Protein Depletion Resin (Thermo Fisher Scientific) was allotted to 1000 µL 96-deepwell plates (Eppendorf). Plasma aliquots were thawed in ice and centrifuged at 13,500 rpm for 5 minutes under 4°C. 10 µL of plasma was loaded into each well and incubated with depletion resin for 30 minutes under room temperature. The solution was filtered by a Multiscreen 96-well PVDF filter plate (Merck Millipore) to collect the depleted sample in a new 1000 µL 96-deepwell plate. 500 µL of cold acetone was added to depleted samples for overnight-precipitation. Protein pellets were collected by 40-minute centrifugation with 2,250g under 4°C, followed by two-round pellet washing with 200 µL cold 95% ethanol.

Purified protein pellets were re-dissolved in 100 µL 2 M Urea/ 25 mM TEABC and incubated on the Eppendorf ThermoMixer at 1,000 rpm for 30 minutes under 29°C. Protein amounts were quantified by Thermo Scientific™ Pierce™ BCA assay kit. 30 µL of 25 mM TCEP/ 25 mM TEABC buffer was added to samples for 30-minute disulfide bond reduction under 29°C. 40 µL of 50 mM IAM/ 25 mM TEABC buffer was added to samples for 30-minute alkylation under 29°C in a dark environment. After the alkylation process, the sample was diluted by adding 30 µL of 25 mM TEABC buffer. For enzymatic digestion trypsin (Promega) and Lys-C (Promega) were added to each sample with the enzyme ratio of 1:20 and 1:40, respectively. After 16-hour digestion, 30 µL of 10% TFA buffer was loaded to each sample to quench the process.

The desalting process was performed with SDB-XC Stage Tip protocol modified from our other study [19]. In brief, the stage tip was packed with 2 layers of SDB-XC membrane cut by a No.16G syringe. The packed stage tips were put on the 96-well adapter to perform centrifugation-based peptide clean-up. The desalting process included five steps: (1) Preconditioning by loading 100 µL buffer B (80% ACN/ 0.1% TFA) with 1,200 g for 3 minutes, (2) Equilibrium by loading 100 µL buffer A (5% ACN/ 0.1% TFA) with 1,200g for 3 minutes, (3)Sample loading with 600g for 10 minutes, (4) Washing by loading 100 µL buffer A with 1,200 g for 5 minutes, and (5) Peptide elution by loading 100 µL buffer B. The peptide samples were collected by loBind 96-well PCR plate (Eppendorf) and quantified by BCA assay. Dried samples were eventually dried and re-dissolved in 2% ACN/ 0.1% FA with 0.0125X spiked-in iRT (Biognosys) before LC-MS/MS analysis.

### LC-MS/MS Analysis

Depleted plasma sample peptides were separated using the Thermo Scientific™ Vanquish™ Neo UHPLC in a trap-and-elute injection configuration and analyzed with the Orbitrap Astral Zoom MS. A Thermo Scientific™ EASY-Spray™ PepMap™ HPLC ES906 column (150 μm x 15 cm, 2 μm pore size) was used for analytical separation. Five hundred nanograms of peptides from individual plasma samples were injected in a randomized order across all sample preparation batches and biological classes. Matrix-matched analytical quality control (QC) pooled peptide samples were run every 30 injections for longitudinal performance monitoring. Experimental sample and analytical QC sample peptides were separated using a 60-sample-per-day (SPD) chromatographic method with a 23.35-minute gradient. Details of the liquid chromatography gradient and liquid chromatography configuration are provided in **Supplemental Table 1**. Eluted peptides were analyzed using narrow window data independent acquisition (nDIA) on the Orbitrap Astral Zoom MS. Parameters for full scan MS1 and nDIA MS2 are listed in **Supplemental Table 2 (A, B, C)**. Six independent gas phase fractionation injections for spectral library assembly were performed on benign, matched control, or cancer plasma peptide pools using the same injection load and gradient. Parameters for gas phase fractionation full scan MS1 and nDIA MS2 are provided in **Supplemental Table 2 (D, E)**.

For unenriched glycopeptide measurements, a random subset of 90 (30 per biological condition) depleted experimental plasma samples were run after the biomarker discovery nDIA study. Peptides were separated and analyzed on the same Vanquish Neo UHPLC in a trap-and-elute injection configuration, coupled to the same Orbitrap Astral Zoom MS, with the same column configuration documented above. Five hundred nanograms of peptides from the subset of individual plasma samples were injected in a randomized block order across all sample preparation batches and biological classes. Each sample block contained a randomized order of 10 benign, 10 healthy, and 10 cancer samples. Matrix-matched analytical QC samples were run every 30 injections for longitudinal performance monitoring. Experimental samples were separated using a 42-minute gradient, whereas analytical QC samples were separated using the same 60SPD LC/MS method and parameters documented above and listed in Tables 1 and 2A-2C. Details of the liquid chromatography gradient and liquid chromatography configuration used for unenriched glycopeptide measurements are provided in **Supplemental Table 3**. Eluted peptides were analyzed using stepped collision energy (SCE) ddMS2 on the Orbitrap Astral Zoom MS. Parameters for full scan MS1 and ddMS2 are listed in **Supplemental Table 4 (A, B, C)**.

### Data Processing

For the canonical plasma proteome search, all mass spectrometer output files were analyzed using Spectronaut v 20.1.250624.92449 (Biognosys AG). A comprehensive spectral library was generated using protein FASTA files against *H. sapiens* (sp_canonical TaxID=9606; 20,433 Swiss-Prot reviewed entries, UniProt). The spectral library was assembled by combining Pulsar search results from individual plasma samples and biological condition pooled plasma peptide gas-phase fractionation runs (3 pools; 1 benign, 1 cancer, 1 control). This combined library was then used to search the experimental data files. Default search parameters were applied during the DIA analysis search. More specifically, enzyme and modification parameters were set at trypsin enzyme specificity with a maximum of two missed cleavages, carbamidomethylation of cysteines (fixed modification), and oxidation of methionine and N-acetylation of proteins (as variable modifications). Spectral matching and identification assignments were performed with a false discovery rate (FDR) of 1% at the precursor and protein group level. Quantification was based on MS2 peak areas, no imputation strategy was used, and local normalization was applied.

For plasma glycoproteome search, the SCE-DDA raw files were imported to Proteome Discoverer v. 2.5 (Thermo Fisher Scientific) using Byonic search engine (Protein Metrics). Human fasta file (v2021-05-06, total 20,324 sequences from human), was used for protein identification. The built-in human N-glycans were integrated from Byonic database, including 132 human N-glycans, 15 human IgM N-glycans, 57 human plasma N-glycans, 59 common biantennary N-glycans and 182 human N-glycans with no multiple fucose were used for the identification of glycan composition. Carbamidomethyl (C) was selected as fixed modification; Acetyl (Protein N-term), deamidation (NQ) and oxidation (M) were selected as common modifications. N-Glycans were selected as “rare”. In the main text, glycans were described as NxHxFxSx: Nx, number(x) of N-acetylglucosamoine (GlcNAc); Hx, number (x) of linked mannose or galactose on antenna; Fx, number (x) of fucose; Sx, number (x) of sialic acids. For example, N4H5F1S2 represents the monofucose-bisialyl-biantennary glycan. The score of identified glycopeptides was higher than 30 and high confidence identification of glycopeptides was set at ≥200; log. probability ≥2; reversed peptide sequence identification was also considered with a protein FDR of 1%, or 20 reverse count. The high confidence of glycopeptide sequence was also set at PEP2D <0.01. The label-free quantitation was further processed by using chromatographic alignment with *m/z* and retention time of identified intact glycopeptides in PD2.5 software.

### Data Analysis

The search results of both proteome and glycoproteome were imported to Perseus v.2.1.5 [20] to conduct the statistical analysis. Proteins below 30% missing value ratio in each group were kept, followed by log_2_ transformation and median subtraction for normalization. Statistical differences among malignancy, benign, and normal groups were first evaluated by one-way ANOVA (p < 0.05). Significant proteins and glycopeptides were further examined using pairwise student’s t-test to determine the cross-group differential abundance. Volcano plots were used to visualize the pairwise expression difference, and the box plots were used to illustrate the differentially expressed proteins across the groups. For the plasma glycoproteomics data, unique glycopeptides were used for analysis to determine the differential site-specific alterations.

## RESULTS AND DISCUSSION

### 96-well Immuno-depletion workflow coupled with the Orbitrap Astral Zoom MS enabled dual-omic profiling in a single-aliquot

Despite the easy accessibility of plasma specimens enabling large-scale proteomic biomarker studies, high-throughput sample preparation and LC-MS/MS with deep proteome coverage, reproducibility and high quantitation accuracy are essential for longitudinal and large-cohort profiling. To address these needs, we developed a single-aliquot, dual-omics workflow that integrates automated 96-well sample processing with ultrafast LC–MS/MS analysis for parallel proteome and glycoproteome profiling at high-throughput. To evaluate the analytical performance of the workflow, we applied it to plasma samples from the Low-Dose Computed Tomography (LDCT) screening cohort, including three participant groups: (i) patients with confirmed lung adenocarcinoma (LUAD), (ii) individuals with pulmonary nodules verified as non-malignant, and (iii) healthy controls (**Figure 1A**).

**Figure 1.**
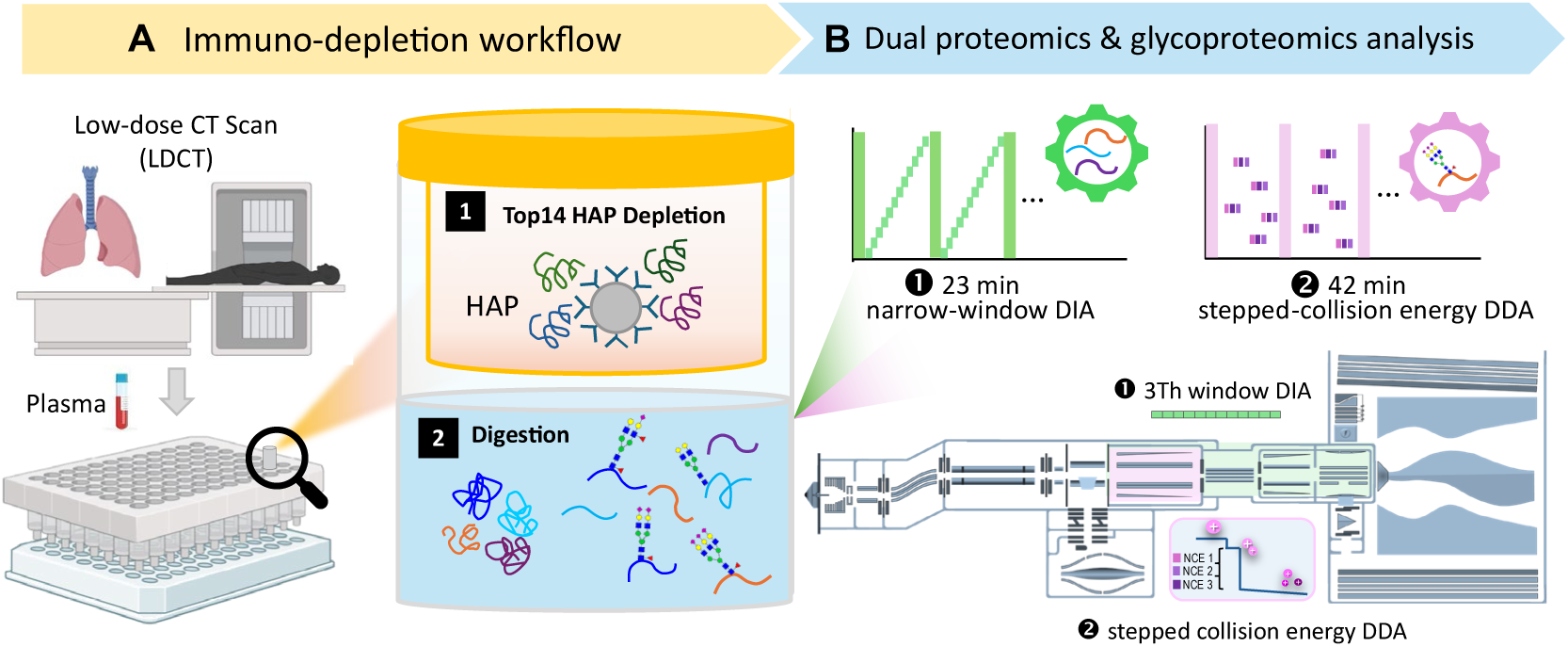
Workflow of concomitant plasma proteomics and glycoproteomics analysis. (A) Plasma samples from individuals with confirmed cancer, benign and normal status after low dose CT screening were analyzed. Sample preparation was performed on 96-well plates. The top 14 highest abundance proteins were first removed by the immunodepletion process, followed by protein digestion. (B) The resulting digests are analyzed by ultrafast LC–MS/MS using Orbitrap Astral Zoom mass spectrometer. Each sample were subjected to two injections for plasma proteomics by narrow window DIA (3Th isolation window) and glycoproteomic profiling by stepped collision energy DDA, respectively.

In the first step, we established a streamlined 96-well plate-based sample preparation that performs immunodepletion of the top 14 most abundant proteins. (**Figure 1A**). The immunodepletion resin, containing antibodies specific to 14 high-abundance plasma proteins (HAPs), such as albumin, immunoglobulin, and apolipoprotein, was allotted into each well for selective capture and removal via subsequent filtration. This step effectively reduced the dominance of HAP concentration to minimize the ion suppression during LC-MS/MS analysis and compressed the dynamic range [21], thereby enhancing detection of mid- and low-abundance proteins. The immunodepletion and subsequent protein digestion are performed in a single automated pipeline with minimal manual intervention, ensuring excellent reproducibility and consistent scalability for hundreds of samples per batch.

After sample preparation, the resulting digests are analyzed by ultrafast LC–MS/MS using Orbitrap Astral Zoom MS. We hypothesize that the superfast acquisition rate (up to 270 Hz) and SCE fragmentation strategy (up to 150 Hz), which alternate HCD collision energies within milliseconds, enhance both peptide and glycopeptide identification efficiency by improving precursor sampling and glycan fragmentation completeness. Thus, two complementary acquisition modes are employed to sequentially analyze both global proteomics and glycoproteomics from the same aliquot: (1) rapid 23-minute data-independent acquisition (DIA) for quantitative proteomics at clinical throughput of 60 samples per day; (2) 42-minute stepped-collision-energy (SCE) data-dependent acquisition (DDA) optimized for glycopeptide fragmentation and identification from unenriched plasma samples.(**Figure 1B**). The workflow enables quantitative dual-omics data acquisition at a rate of ∼30 patients per day, providing a practical foundation for longitudinal studies of biomarker discovery. Notably, both analyses originated from the same aliquot of the immunodepleted plasma digest, eliminating the need for independent sample preparation for different omic studies. This strategy ensured identical sample composition between proteome and glycoproteome workflows, thereby reducing the sample preparation bias, improving data comparability and minimizing variability arising from separate preparation batches. The single-aliquot-dual-acquisition approach achieved characterization of both protein abundance and glycosylation features from the same biological specimen, maximizing analytical throughput and sample utilization efficiency.

### Establishment of a multi-layer quality control system for high-throughput plasma analysis

Robust quality control (QC) is essential for ensuring analytical reproducibility and data comparability in high-throughput LC–MS–based plasma proteomics. To evaluate the robustness and temporal stability of the established workflow, we implemented a three-layer QC strategy to monitor performance at the instrument, analytical, and sample preparation levels. Three types of quality control samples were prepared and inserted in the data acquisition queue (**Figure 2A**), including (1) instrument QC (system performance monitoring): Instrumental stability was assessed daily using a standard HeLa tryptic digest as a system suitability test in chromatographic retention time stability, MS1 and MS2 signal intensity, and identification rates; (2) matrix-matched plasma QC (analytical process control): the overall stability of LC–MS performance for complex plasma matrices across every 30 analytical runs was evaluated by a pooled depleted plasma reference sample from the study cohort; (3) plate QC (sample preparation reproducibility): a designated plate QC sample from depleted plasma pool was included in each batch to monitor intra- and inter-plate variability introduced during immunodepletion, digestion, and cleanup in the 96-well plate-based sample preparation.

**Figure 2.**
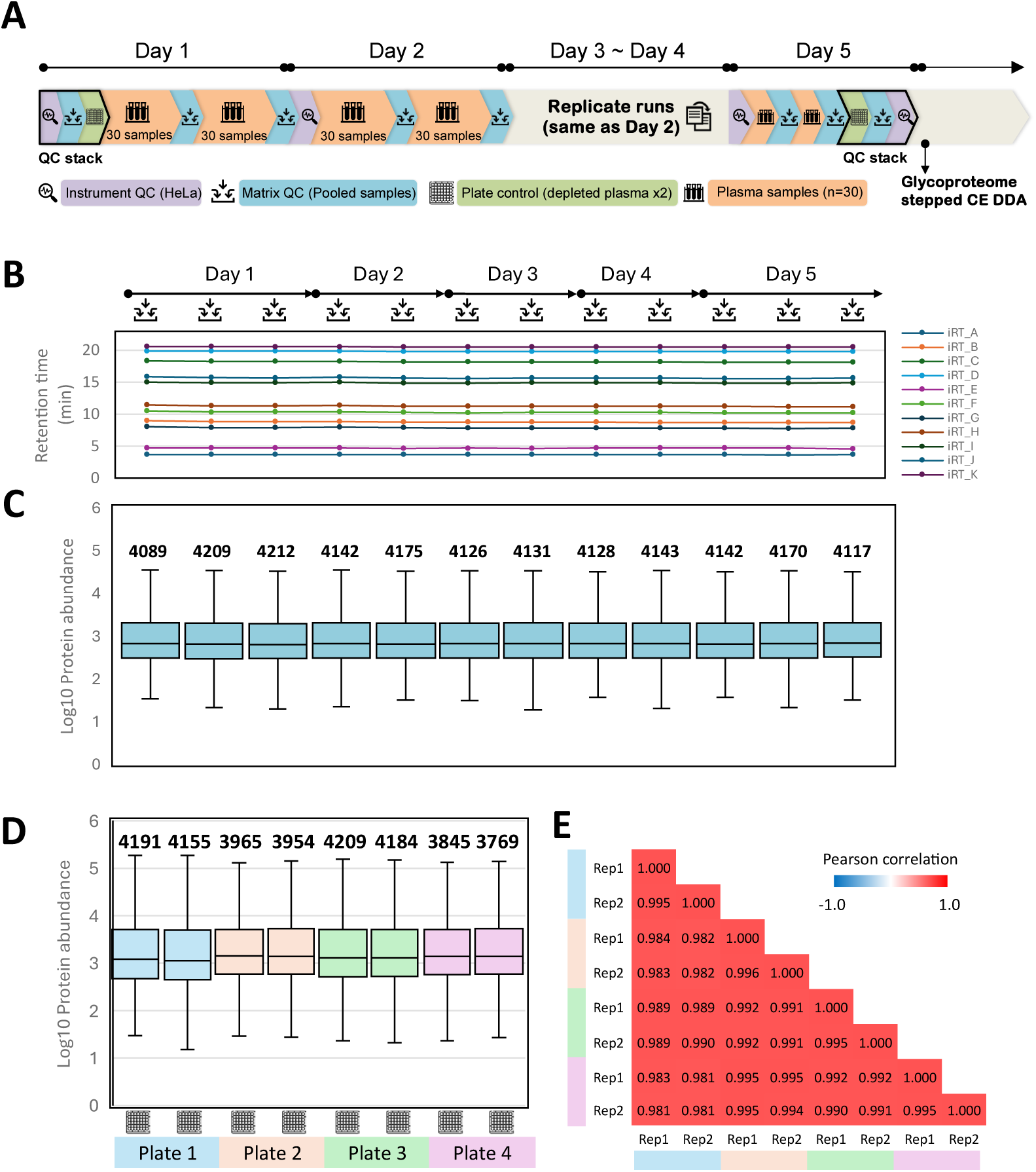
Monitoring the instrument robustness and plate reproducibility with quality control and plate reference samples. **(A)** Three types of quality control (QC)samples were prepared and inserted in the data acquisition queue, including instrument QC (HeLa cells), matrix QC (pooled plasma sample), and plate QC (depleted plasma sample). To monitor the longitudinal instrument status, instrument QC was analyzed daily, and matric QC was analyzed after every 30 sample injections throughout the study. **(B)** Reproducible retention time profile of 11 spiked-in empirical indexed retention time (iRT) peptides across 14 matrix QC samples. **(C)** Number of identified protein groups and protein abundance distribution throughout the 14 matrix QC samples. **(D)** Number of identified protein groups and protein abundance distribution of technical duplicates of plate QC samples across four 96-well plates. **(E)** The Pearson correlation coefficient between technical duplicates of plate QC samples across four 96-well plates.

Across all QC layers, the workflow demonstrated excellent stability and reproducibility across 5-days of data acquisition. The retention time profiling of 12 spiked-in iRT peptides from plasma pool QC samples were monitored and used to track the chromatographic alignment throughout the whole study (**Figure 2B**). A highly reproducible retention time was observed with < 8% CV%, confirming the robustness of the short 23-min gradient for long-term continuous analysis. Furthermore, consistent identification numbers were observed with an average of 4,149 ± 36 protein groups (PGs) and similar intensity distribution spanning over 6 orders of magnitude (**Figure 2C**). Plate QC samples processed independently across 96-well plates also showed minimal variation in protein identification numbers (4,034 ± 173) and quantitative profiles (3,935 proteins groups below 10%CV with ≥ 60% quantification coverage) (**Figure 2D**), indicating reliable depletion and digestion efficiency. This result demonstrated the feasibility of the larger-scale sample preparation workflow, as it shows excellent performance at a rate that exceeded the speed of the MS instrument.

### Deeper plasma proteome profiling enhanced the potential cancer biomarker screening

With the acquisition throughput of 60 samples per day and the added depth provided by the sample pool specific gas-phase-fractionation spectral library, the global proteomics analysis achieved the depth of average identification of 3,756 PGs (±11.0%) per sample (**Figure 3A**). The total number identified was over 4,900 PGsacross 300 individuals, which surpassed the depth reported in previous immunodepletion-based studies and represented one of the most comprehensive proteome coverage achieved so far [14,21,22,23]. Compared to the latest version of Human Plasma PeptideAtlas (2025-8), identified 1,599 previously unseen proteins [24] (**Figure 3B**). Compared with the recent benchmarking studies evaluating several plasma proteomic workflows, including PEA, depletion, and nanoparticle-based enrichment [14], our method demonstrated comparable or superior analytical depth. Compared to nanoparticle-based enrichment approach, immunodepletion-based approaches require substantially less plasma input, which highlighted the efficiency and clinical applicability for limited-volume cases.

**Figure 3.**
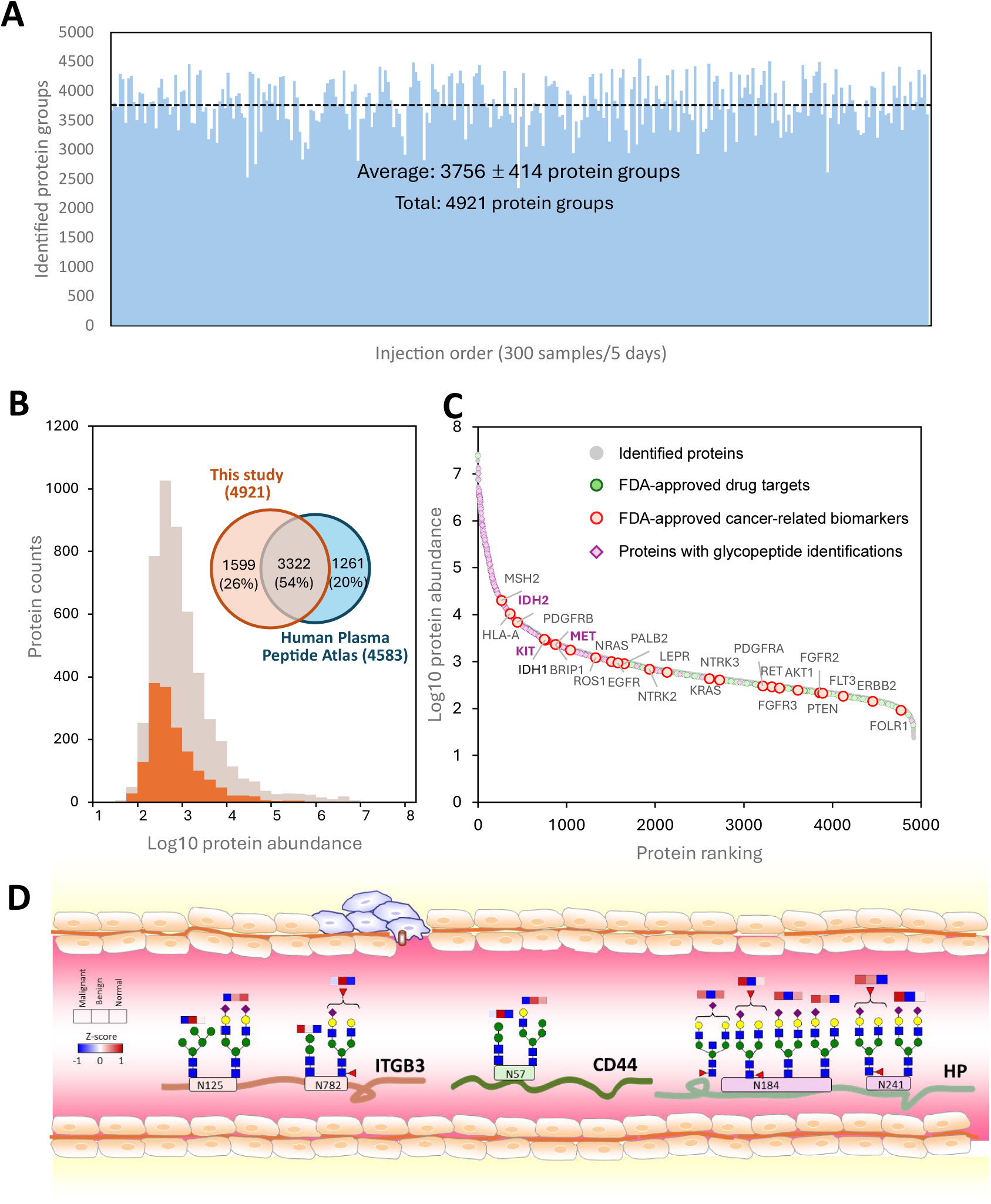
Summary of protein identification coverage of 300 plasma samples analyzed in 5 days (60 samples per day). **(A)** Overview of identified protein groups across 300 samples, sorted by the order of acquisition sequence. **(B)** The comparison between the identification results and the Human Plasma Peptide Atlas database. The histogram represents protein intensity distribution of the identification result, where the proteins not reported in the Human Plasma Peptide Atlas are highlighted. **(C)** The dynamic range of protein abundance of 4921 identified proteins. Green data points represent proteins reported as FDA-approved drug targets. Red data points represent the FDA-approved cancer-related biomarkers. Proteins identified with glycopeptides were highlighted with purple diamonds or bold purple fonts. **(D)** Representative glycoproteins identified from the samples.

The protein abundance distribution revealed a dynamic range spanning over 6 orders of magnitude (**Figure 3C**). Among the 4,921 proteins, 303 proteins had been reported with differential expression in cancerous tumors in Human Protein Atlas (HPA, https://www.proteinatlas.org/). 24 proteins were FDA-approved cancer drug targets or biomarkers, where 7 proteins were lung cancer therapeutic biomarkers, such as EGFR, KRAS, and ERBB2 in the low abundance region. Other 13 proteins are cancer biomarker candidates in investigated phase reported in MarkerDB [25]. The detection of these clinically approved or under-investigated biomarker or drug targets demonstrated the high sensitivity for biomarker studies.

For glycoproteome profiling, a total of 454 glycoproteins were identified. The majority are located in the higher-abundance region, while several glycopeptides were also identified from low-abundance proteins (**Figure 3C**). The coverage also includes three clinical biomarkers (IDH2, KIT, and MET) and 74 FDA-approved drug targets. To illustrate representative site-specific glycosylation patterns across clinical cohorts, selected glycopeptides from ITGB3, CD44, and HP, along with their corresponding N-glycosylation sites, are shown in **Figure 3D**. The results reveal differential degree among multiple site-specific glycosylation in the three groups. Among cancer-associated up-regulation, HP, an acute-phase protein, demonstrates site-dependent glycan alterations, where N184 and N241 exhibit enhanced abundance of multi-antennary sialylated and fucosylated structures in malignant samples, aligning with reported cancer-associated glycoproteins [26,27,28,29]. ITGB3 glycopeptides at N782 exhibit enhanced sialylation and fucosylation glycan in benign individuals. Interestingly, CD44, a known metastasis-associated receptor [30,31], also shows elevated complex-type glycan abundance at N57 in the benign individuals. Together, these results highlight disease-specific glycosylation regulation and emphasize the biological relevance of site-specific glycoproteome, supporting the utility of glycoform-level biomarkers in cancer detection and characterization.

### Comparative evaluation for inter-instrument transferability

Data consistency across evolving instrumentation platforms is an essential requirement for long-term clinical studies, biobanking efforts, and inter-laboratory comparisons. To ensure such transferability, we evaluated the performance and compatibility of our high-throughput plasma proteomics workflow on two generations of Orbitrap mass spectrometers by 56 samples following Top14 protein depletion. Using a 120-minute gradient on the Orbitrap Fusion Lumos MS, a total of 2,084 PGswere quantified at a rate of 17.4 PGs per minute (**Figure 4A**). In contrast, the Orbitrap Astral Zoom MS achieved 112.7 PGs per minute, representing a 4.4 to 9.8-fold increase in profiling speed while maintaining greater proteome depth (**Figure 4B**). This throughput also exceeds recent reports using similar Orbitrap Astral MS setup (e.g., 1,307 PGs in 24 min ≈ 54 PGs/min). The faster acquisition speed also enhances the number of peptides for quantification, 11.1 peptides/protein on the Orbitrap Astral Zoom MS versus 8.8 peptides/protein on the Orbitrap Fusion Lumos MS. Accordingly, the protein quantitation coverage was also improved using the Obritrap Astral Zoom MS, quantifying 78% proteins across 80% samples, which is 13% increase of proteins compared to Orbitrap Fusion Lumos MS results (**Figure 4A**). Nevertheless, the quantified protein abundances show high correlation coefficient exceeding 0.90 across the two platforms, confirming strong reproducibility and supporting seamless integration of datasets across instrument generations or inter-instrument transferability (**Figure 4C**). These results validate the robustness and scalability of our workflow for longitudinal and multi-center plasma proteomics studies.

**Figure 4.**
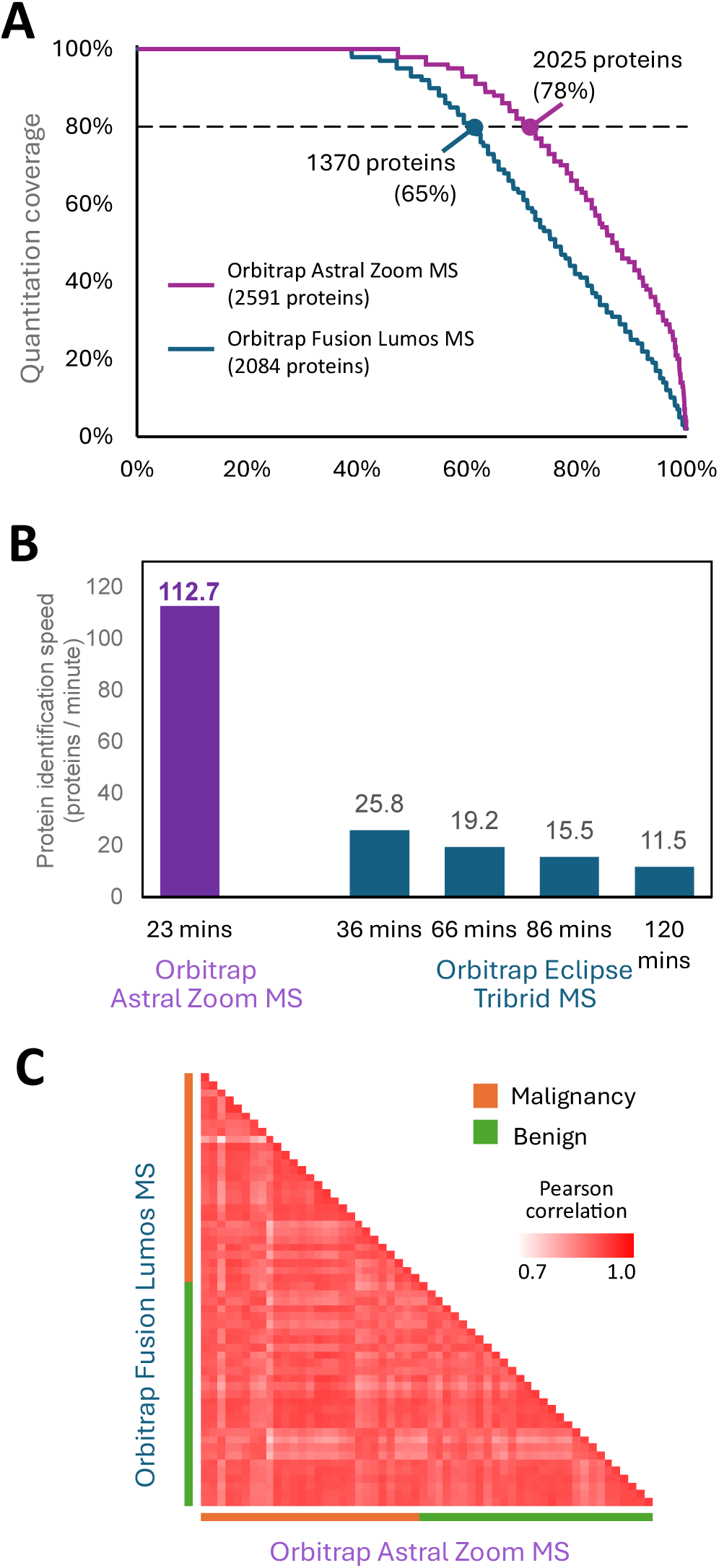
The benchmark between 60-SPD plasma proteomic pipeline with other longer-gradient methods using other MS. **(A)** The quantitation coverage comparison between Orbitrap Astral Zoom MS and Orbitrap Fusion Lumos MS, where the dashed line represents the 80% quantitation coverage. **(B)** The protein profiling speed comparison between Orbitrap Astral Zoom MS and Orbitrap Eclipse Tribrid MS using 3 different analytical throughputs. The speed is calculated by the number of identified proteins divided by the gradient time. **(C)** The heatmap of the inter-sample Pearson correlation coefficient of protein intensities acquired from the same samples using Orbitrap Fusion Lumos and Orbitrap Astral Zoom MS

### Enrichment-free glycoproteome: strong MS/MS evidence and high-confidence glycopeptide identification

To evaluate the analytical performance of enrichment-free plasma glycoproteomic analysis, a direct comparison was performed with the conventional enrichment-based workflow using two sets of identical paired plasma samples: peptide samples without enrichment (n=3) and glycopeptides after ZIC-cHILIC enrichment and analyzed by stepped collision energy (SCE)-DDA mode. **Figure 5A** revealed that the majority of glycopeptides are highly confident (Byonic score >200) in both enriched (92%) and enrichment-free (78–84%) paired samples. The higher proportion of low-confidence identification in the enrichment-free sample compared to the enriched sample is expected, given the increased sample complexity in the absence of enrichment. Nevertheless, the enrichment-free workflow still achieved a majority of high-confidence glycopeptides (≥78%), demonstrating that reliable identification can be obtained without prior enrichment. High-confidence glycopeptides were detected in both enriched (92%) and enrichment-free (78–84%) samples.

**Figure 5.**
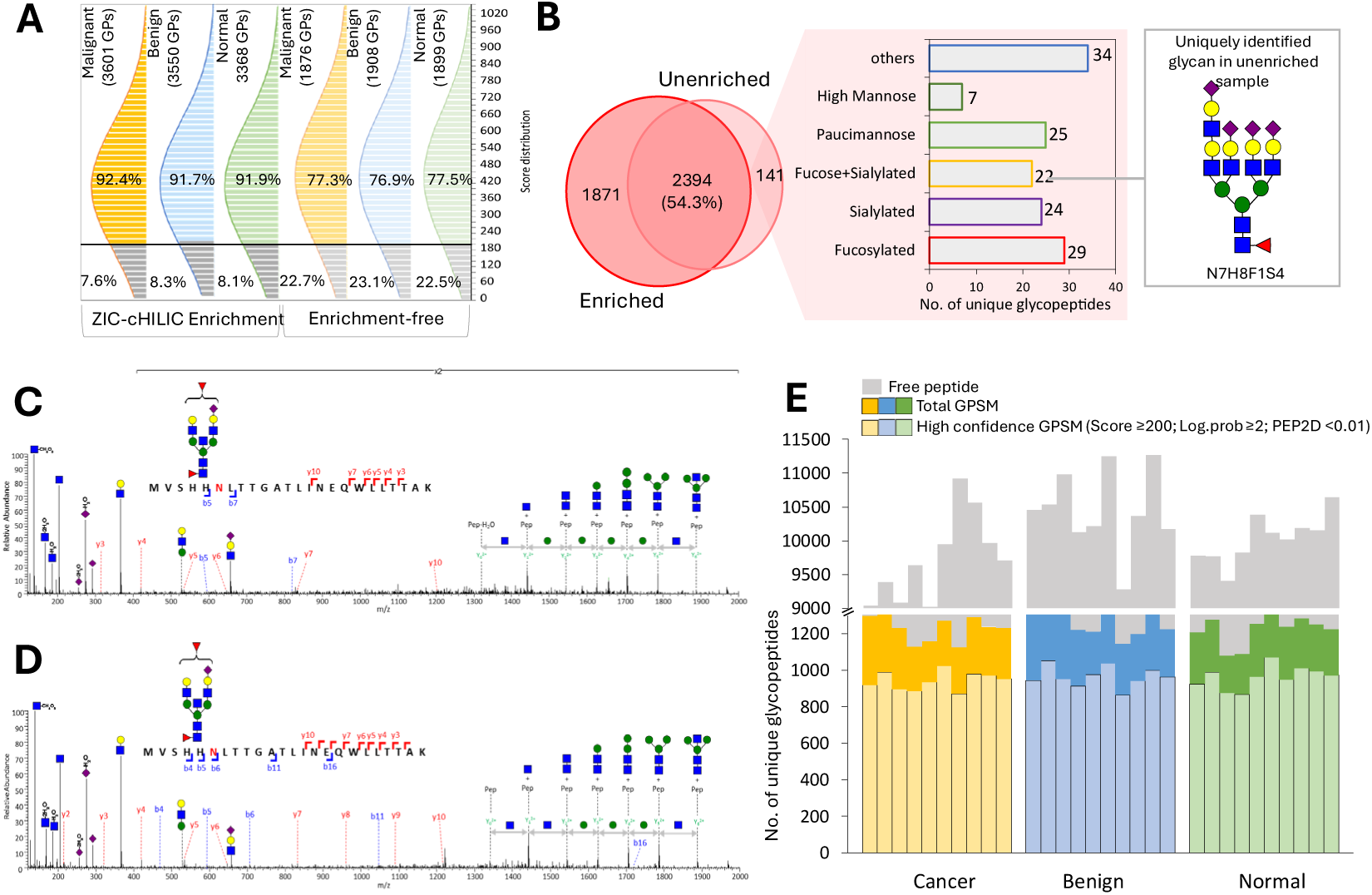
Summary of enrichment–free glycoproteomic analysis across malignant, benign and normal group. To evaluate the performance of enrichment-free analysis, comparative analysis was performed for depleted samples without enrichment (enrichment-free) and enrichment with ZIC-cHILIC. **(A)** Byonic score distribution (Score ≥200) **(B)** Venn diagram of unique glycopeptide overlap between enrichment-free and enriched sample. Comparison of identical N-glycopeptides MS/MS spectra from Hp glycoprotein with N5H5F2S1 glycan were observed from **(C)** enriched sample (*m/z* 122.0497, z = 4+, Byonic score = 206.8) and **(D)** from enrichment-free sample (*m/z* 1222.0526, z = 4+, Byonic score = 547.9). Summary of enrichment-free glycoproteomic analysis across groups (*n=10* for each group) using Stepped Collision Energy (SCE) mode with DDA acquisition. **(E)** Number of total and high confidence (Byonic score ≥200, log. Probability ≥2 and PEP2D <0.01) glycopeptides across clinical group. Blue squares, GlcNAc; green circles, Man; yellow circles, Gal; red triangles, fucose; purple diamond, sialic acid.

A glycopeptide distribution map further illustrated that both enriched and enrichment-free samples exhibited highly similar chromatographic and mass-to-charge distribution (**Figure S1**). In both cases, glycopeptides were primarily detected across the 800-1600 m/z range and equally distributed throughout the 40-minute retention time, confirming that the overall ionization and chromatographic features of detected glycopeptides remained consistent between methods. The only observable difference was the expected higher number of identified glycopeptides in enriched (4265 glycopeptides) compared to enrichment-free (2535 glycopeptides) samples. The overlap of identified glycopeptides between enriched and enrichment-free workflows was assessed using a Venn diagram (**Figure 5B**). In total, 54.3% (2394 glycopeptides) were shared between the two datasets, with 1871 and 141 glycopeptides being uniquely identified in enriched and enrichment-free samples, respectively. Notably, these 141 unique glycopeptides consist of diverse glycan types: Fucosylated (29), sialylated (24), fuscose+sialylated (22), paucimannose (25), high-mannose (7), and others (34) glycopeptides. In addition, a particularly notable glycan, N7H8F1S4, was exclusively observed in the enrichment-free sample. This tetra-antennary, fucosylated and highly sialylated glycan type has been widely associated with tumor progression, cancer cell migration and metastasis [32]. These results suggest that the enrichment-free workflow may capture glycopeptides that are lost or selectively excluded during enrichment, which supports the idea that the enrichment-free workflow can complement enrichment-based methods and expand the overall glycopeptide landscape.

To further evaluate the reliability of enrichment-free identifications, MS/MS spectra of two identical N-glycopeptides derived from haptoglobin (Hp) carrying the N5H5F2S1 glycan composition were compared between enriched and enrichment-free analyses (**Figure 5C– D**). Both spectra showed similar fragmentation patterns, including distinct *b/y-ions* (b4-b7, b11, b16, and y2-y10) from the peptide backbone and diagnostic glycan fragments such as oxonium ions (*m/z* 366.14 for HexHexNAc+, 204.09 for HexNAc+, 138.06 for HexNAc-CH_6_O_3_, 168.07 for HexNac-2H_2_O, 274.09 for Neu5Ac-H_2_O+, and 292.09 for Neu5Ac+) and Y-series ions, including Y0 (naked peptide), Y1 (peptide+HexNAc), Y2 (peptide+2HexNAc), Y3 (peptide+2HexNAcHex), Y4 (peptide+2HexNAc2Hex), Y5 (peptide+2HexNAc3Hex), and Y6 (peptide+2HexNAc4Hex). The close resemblance of fragmentation profiles confirms that MS/MS spectra quality is not compromised by the absence of enrichment. These findings confirm that the enrichment-free workflow provides sufficient sensitivity and spectral quality to ensure confident glycopeptide identification. Although a modest drop in peptide identification number and confidence is unavoidable, it does not critically compromise the overall data quality.

### Evaluation of enrichment-free glycoproteomics workflow across clinical cohorts

The enrichment-free glycoproteomic workflow was next evaluated for performance across malignant (*n = 10*), benign (*n = 10*), and normal (*n = 10*) plasma samples collected over a 42-minute gradient using stepped collision energy (SCE)-DDA (**Figure 5E**). Across all samples, the total number of identified glycopeptides was consistent, ranging approximately 1100– 1300 per run, with 800-1000 classified as high-confidence identifications (Byonic score ≥200, log. probability ≥2, PEP2D <0.01). Notably, more than 82% of all glycopeptides in each group met the high-confidence criteria (**Figure S2.A-C**), suggesting the reliability and reproducibility of the enrichment-free workflow.

To evaluate the glycan and glycosite microheterogeneity, the number of unique glycans and N-glycosites per glycoprotein was compared across clinical cohorts. For glycan micro-heterogeneity (**Figure S3.A**), approximately 250 glycoproteins were identified with 1-5 unique glycans, with single-glycan occupancy being the most commonly detected (normal = 106, benign = 107, and malignancy = 105 glycoproteins). Protein decorated with two or more distinct glycans occurred less frequently, and their proportions were comparable across the clinical cohorts. Similarly, glycosite heterogeneity (**Figure S3.B**) followed the same pattern, with most glycoproteins containing a single N-glycosite (normal = 161, benign = 195, malignant = 173). Multi-site glycoproteins represented a smaller fraction, again showing minimal variation between groups. The glycan type distribution across clinical cohorts show major classes in sialylation (32%) and complex fucose+sialylation (28%) compared to fucosylation (13%), paucimannose (7%), high mannose (6%), and others (15%) (**Figure S3.C)**. The trends remained similar among malignant, benign, and normal cohorts. Although the parallel trends in identification results did not show distinct differences between cohorts, it is important to note that these results represent identification-level frequencies rather than quantitative abundance. The biologically meaningful differences in relative expression abundance or site-specific occupancy will be derived by further quantitative and statistical analyses.

Collectively, these findings suggest that the enrichment-free workflow preserves both glycan and glycsosite detectability with stable and reproducible identification coverage across diverse plasma samples while maintaining a high proportion of confident glycopeptide assignments. This high analytical consistency, achieved without enrichment, highlights the robustness and practicality of the workflow for large-scale or clinical glycoproteomic studies.

### Integrated plasma proteome and glycoproteome profiling reveals complementary biomarker insights

Our dual-omic strategy enables simultaneous assessment of changes in protein abundance and glycosylation patterns, thereby broadening the detection space for potential biomarkers. Parallel statistical testing was performed to explore differential proteins and glycopeptides across cohorts. An initial one-way ANOVA test discovered 186 proteins and 47 glycopeptides showing statistical significance in abundance (p < 0.05). The heatmap and a full list of 47 significant glycopeptides can be found in **Figure S4** and **Supplemental Table 5**, respectively. These candidates were further examined using pairwise Student’s *t*-tests and visualized as volcano plots.

The pilot study aims to identify biomarkers to differentiate cancer from benign groups. Between malignancy and benign groups (**Figure 6A**), the volcano plot reveals many proteins upregulated in the cancer group, such as several S100 proteins (S100P, S100A8, S100A9, and S100A12) and annexin family proteins (ANXA1 and ANXA6), which have been reported to regulate cancer cell proliferation, apoptosis, and tumor microenvironment development [33,34]. We further evaluated the expression profiles of 46 proteins showing statistical significance specific to malignant cases (**Figure 6B**). Among these, ARHGEF1 and MYCBP were markedly upregulated in the malignancy group. ARHGEF1 has recently been linked to immune modulation [35], while MYCBP (AMY-1), a co-activator of c-MYC, has been reported as highly expressed in several tumor types, including basal-like breast cancer [36,37]. Conversely, LOX1 and MYOC exhibited lower abundance in malignancy samples. LOX1 downregulation aligns with Human Protein Atlas (HPA) data showing reduced expression in LUAD tumors [38], while MYOC reduction is consistent with its loss in cancer-associated cachexia [39].

**Figure 6.**
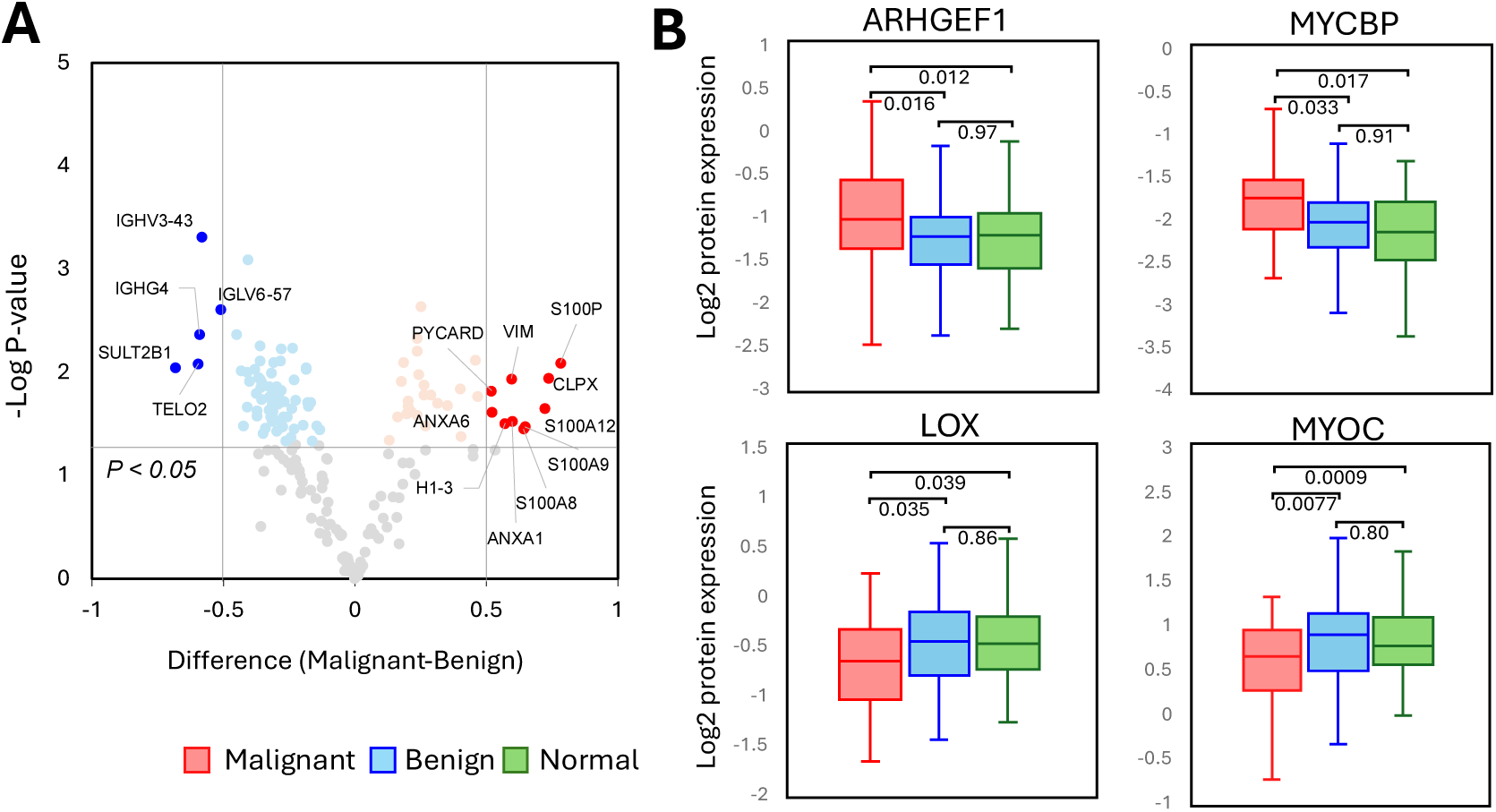
Quantitative proteomic analysis across clinical groups. (A) Volcano plot of significant differential proteins between malignant and benign groups. (B) Selected examples of differentially expressed proteins across malignant, benign and normal groups

At the site-specific glycopeptide levels (**Figure 7A**), 13 and 24 glycopeptides were found to be up-regulated and down-regulated in malignancy across the malignant group compared to either benign or normal groups, respectively. Notably, glycosylation of KNG1 at N294 (N4H5F2S1) exhibits consistent up-regulation, whereas 6 (FN1_N528_N4H5S1, C4A_N226_N6H3, CNDP1_N382_N5H6S3, PROC_N290_N4H5S2, CLTC_N720_N2, PI16_N403/N409_N1F1) were consistently down-regulated in both malignant-benign and malignant-normal pairwise comparisons. In the benign-normal comparison, 7 glycopeptides exhibited consistent up-regulation, and 2 showed consistent downregulation in the benign group when compared with both malignant and normal samples. These consistent upregulation and downregulation across independent pairwise comparisons nominate their potential as site-specific glycopeptide biomarkers associated with malignant and benign groups. In addition, no changes were observed at the corresponding protein level for these glycopeptides, suggesting that the observed difference arises from site-specific glycosylation regulation rather than altered protein abundance. This underscores the importance of glycosylation profiling as a complementary insight into disease biomarker discovery.

**Figure 7.**
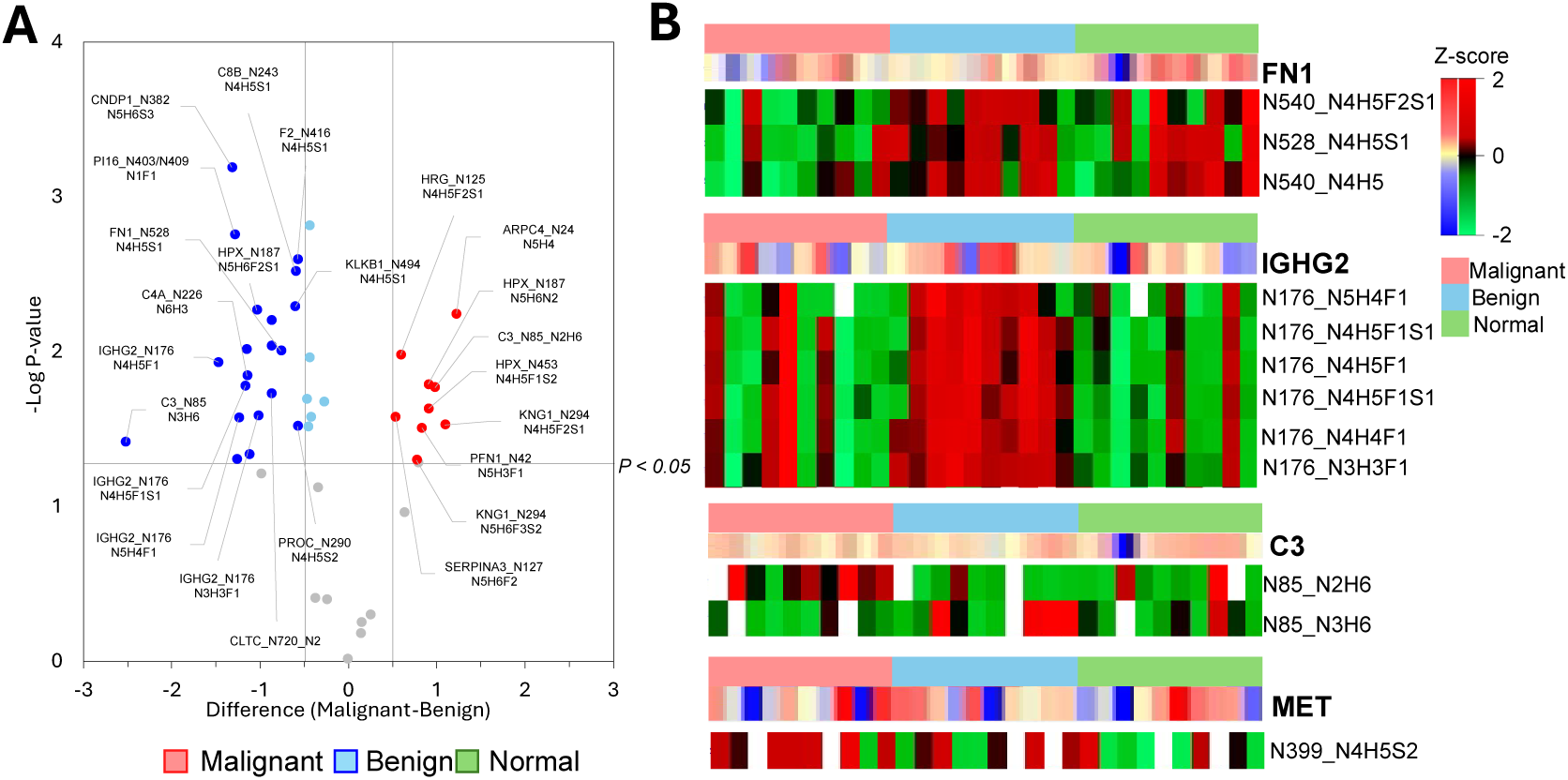
Quantitative glycoproteomic analysis across clinical groups. (A) Volcano plot of significant differential glycopeptides between malignant and benign groups. (B) Selected examples of differentially expressed glycopeptides across malignant, benign and normal groups

Finally, we illustrate selected examples for the relationship between personalized protein expression and corresponding glycopeptides patterns across the clinical cohort (**Figure 7B**). Across all four representative glycoproteins (FN1, IGHG2, C3, and MET), no significant differences were observed at the protein level, yet site-specific glycosylation displayed distinct regulation at glycopeptide level. Specifically, upregulation of a few glycopeptides were observed in C3 (N85_N3H6) and MET (N399_N4H5S2) from malignant cases. In IGHG2, most of the glycopeptides exhibited elevated abundance predominantly within the benign cohort, which shows more dramatic change compared to the protein level. In FN1, the glycopeptide mapped to FN1 showed down-regulation in the malignant group, though a few cases show higher protein abundance, compared to the benign and normal groups. These results also emphasize that glycosylation changes can occur independently of total protein abundance, highlighting the biological importance of site-specific glycosylation as an additional layer of molecular regulation potentially relevant to disease differentiation and biomarker discovery.

## CONCLUSION

Our single-aliquot, enrichment-free dual-omics pipeline provides a scalable and reproducible platform for simultaneous plasma proteome and glycoproteome profiling. Taking advantage of high speed in both data acquisition for DIA and SCE-DDA, deep proteome and glycoproteome coverages were achieved from the same sample at 24 samples per day. The integrated multi-layer quality control system verified the robustness of sample processing, column separation, and instrument stability over extended multi-day acquisitions, which supports reliable operation in large-scale clinical studies. This workflow accelerated plasma protein profiling speed while maintaining comprehensive depth, identifying nearly 5000proteins, including 1599 proteins absent from the Human Plasma PeptideAtlas, thereby expanding the detectable proteome space. The strong inter-instrument protein abundance correlation (r > 0.9) further demonstrates cross-platform transferability and reproducibility, highlighting the method’s analytical robustness. As a proof of concept, differential expression analysis in plasma from lung cancer vs nodule cohorts highlighted several candidate biomarkers, such as S100 and annexin family proteins, that warrant further validation in ongoing studies.

Without enrichment, our platform provides high-confidence glycopeptide identification (≥78%) with MS/MS spectrum quality with higher similarity to enriched counterparts, which eventually identified comparable glycan distribution and glycosite number per protein across clinical cohorts. In addition, the identification of a unique tetra-antennary fucosylated and multi-sialylated glycan (N7H8F1S4) exclusively in the enrichment-free sample highlights the workflow’s ability to capture cancer-associated glycoforms without enrichment steps. Quantitative analysis across normal, benign, and malignant cohorts revealed that site-specific glycosylation changes provide meaningful insight into disease-associated regulation, even in the absence of protein level changes. Representative examples such as FN1, IGHG2, C3, and MET also demonstrated distinct glycopeptide-level changes without corresponding protein-level changes, suggesting that site-specific glycosylation regulation operates as an independent regulatory layer impacting protein function, cell-signaling, cell adhesion, cell migration and invasion. These results demonstrate that the enrichment-free workflow achieves comparable depth, fragmentation quality, and reproducibility to conventional enrichment-based approaches, offering a robust alternative for large-scale plasma glycoproteomic studies, offering insight that site-specific glycosylation unravels subtle yet biologically relevant disease signatures that conventional proteomic analysis may overlook.

Overall, this dual-omics platform establishes a technically robust and high-throughput framework for clinical plasma proteomics, which offers a foundation for scalable biomarker discovery and multi-omics integration in translational research.

## ACKNOWLEDGMENT

This study was supported by National Science and Technology Council (NSTC 113-2113-M-001-020-MY3), Key and Novel Therapeutics Development Program for Major Diseases (ASKPQ-111-KNT) and Academia Sinica (AS-GC-111-M03) in Taiwan. The instrument analysis and partial reagents were supported by Thermo Fisher Scientific.

## Notes

### Competing Interest Statement

The instrument analysis and partial reagents in this study were supported by Thermo Fisher Scientific.

### Summary of Updates

Revision for the Introduction section: (1) Removing the repeating sentence (2) Revising the literature review of Oxoscan-MS

