## Supplemental Table S2 for "A Single-Aliquot, Enrichment-Free Workflow for High-Throughput Plasma Proteome and N-Glycoproteome Profiling"

**Table S2.** Orbitrap Astral Zoom mass spectrometer parameters. (A) Global source and mass spectrometer parameters; (B) MS1 full scan experiment parameters; (C) MS2 DIA scan experiment parameters; (D) Gas phase fractionation MS1 full scan experiment parameters; (E) Gas phase fractionation MS2 DIA scan experiment parameters.

**A**

| Global Parameters (Source & MS). | |
|---|---|
| Positive Ion Voltage | 2100 Volts |
| Ion Transfer Tube Temperature | 290°C |
| Expected Peak Width | 6 seconds |
| Default Charge State | 2 |
| Lock Mass Correction | Off |

**B**

| MS1 Full Scan Experiment Parameters | |
|---|---|
| Orbitrap Resolution | 240K |
| Scan Range (m/z) | 380-980 |
| RF Lens (%) | 40 |
| Normalized AGC Target (%) / Absolute AGC Value | 500% / 5.00e6 |
| Maximum Injection Time | 5 milliseconds |
| Microscans | 1 |

**C**

| MS2 DIA Scan Experiment Parameters | |
|---|---|
| Precursor Mass Range (m/z) | 380-980 |
| Isolation Window (m/z) | 3 |
| Window Placement Optimization | On |
| AGC Target | Custom |
| Normalized AGC Target (%) / Absolute AGC Value | 500% / 5.00e4 |
| Maximum Injection Time | 7 milliseconds |
| DIA Scan Range (m/z) | 150-2000 |
| HCD Collision Energy (%) | 25 |
| RF Lens (%) | 40 |
| Pre-Accumulation | On |
| Loop Control | Time |
| Time | 0.6 seconds |

**D**

| GPF MS1 Full Scan Experiment Parameters | |
|---|---|
| Orbitrap Resolution | 240K |
| Scan Range (m/z) | Incremental 100 m/z scans (380-480; 480-580; 580-680; 680-780; 780-880; 880-980) |
| RF Lens (%) | 40 |
| Normalized AGC Target (%) / Absolute AGC Value | 500% / 5.00e6 |
| Maximum Injection Time | 3 milliseconds |
| Microscans | 1 |

**E**

| GPF MS2 DIA Scan Experiment Parameters | |
|---|---|
| Precursor Mass Range (m/z) | Incremental 100 m/z scans (380-480; 480-580; 580-680; 680-780; 780-880; 880-980) |
| Isolation Window (m/z) | 1 |
| Window Placement Optimization | On |
| AGC Target | Custom |
| Normalized AGC Target (%) / Absolute AGC Value | 500% / 5.00e4 |
| Maximum Injection Time | 18 milliseconds |
| DIA Scan Range (m/z) | 150-2000 |
| HCD Collision Energy (%) | 25 |
| RF Lens (%) | 40 |
| Pre-Accumulation | On |
| Loop Control | Time |
| Time | 0.6 seconds |
