## Supplemental Table S1 for "A Single-Aliquot, Enrichment-Free Workflow for High-Throughput Plasma Proteome and N-Glycoproteome Profiling"

**Table S1.** Gradient specifications and LC Configuration of narrow-window DIA for plasma proteome

| 60 SPD Gradient Specifications and LC Configuration | | | |
|---|---|---|---|
| Gradient | Time (min) | % Mobile Phase B | Flow (µl/min) |
| | 0 | 10 | 2.0 |
| | 0.3 | 10 | 2.0 |
| | 0.6 | 10 | 0.8 |
| | 13.6 | 22.5 | 0.8 |
| | 20.5 | 35.0 | 0.8 |
| | 20.9 | 55.0 | 2.0 |
| | 20.95 | 99.0 | 2.0 |
| | 22.35 | 99.0 | 2.0 |
| LC Parameters | LC Configuration | Trap and Elute | |
| | Fast Loading/Equilibration Mode | Pressure Control | |
| | Loading/Equilibration /Wash Pressure | Max Pressure | |
| | Equilibration Factor | 3 | |
| | Sampler Temperature | 7°C | |
| | Mobile Phase A / Weak Wash | 0.1% Formic Acid in Water | |
| | Mobile Phase B / Strong Wash | 0.1% Formic Acid in 80% Acetonitrile | |
| | Zebra Wash | Enabled | |
| | Zebra Wash Cycles | 4 | |
| | Analytical Column Temperature | 50°C | |
| Column Specifications | Analytical Column | EASY-Spray™ PepMap™ Column, 2µm C18, 150µm × 15 cm (P/N ES906) | |
| | Trap Column | PepMap™ Neo Trap Cartridge, 5 µm C18 300 µm x 5 mm, (P/N 174500) | |
