## Supplementary material for "A Single-Aliquot, Enrichment-Free Workflow for High-Throughput Plasma Proteome and N-Glycoproteome Profiling": Figure S4

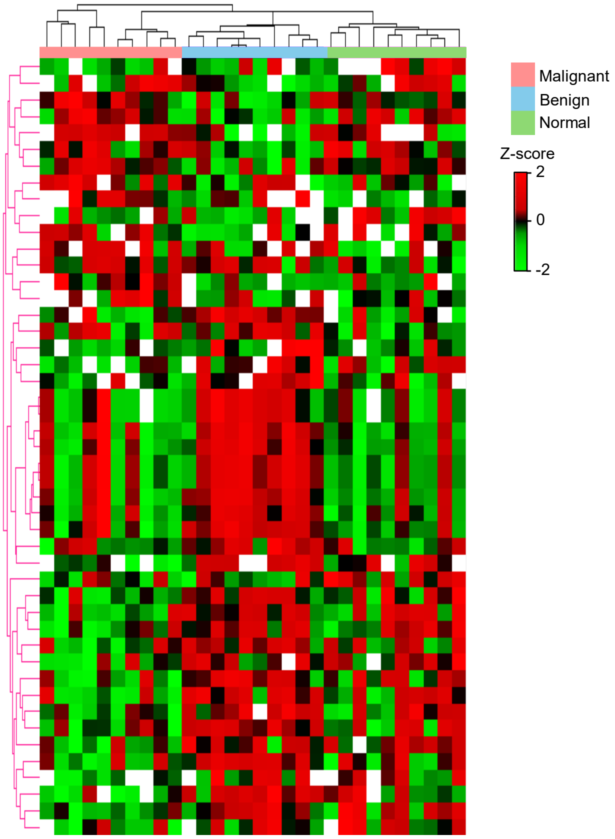


**Figure S4.** Heatmap of 47 significant (*p-value* <0.05) glycopeptide expressions across clinical group (*cancer n=10, benign n=10, normal n=10*).
