## Supplementary material for "A Single-Aliquot, Enrichment-Free Workflow for High-Throughput Plasma Proteome and N-Glycoproteome Profiling": Figure S3

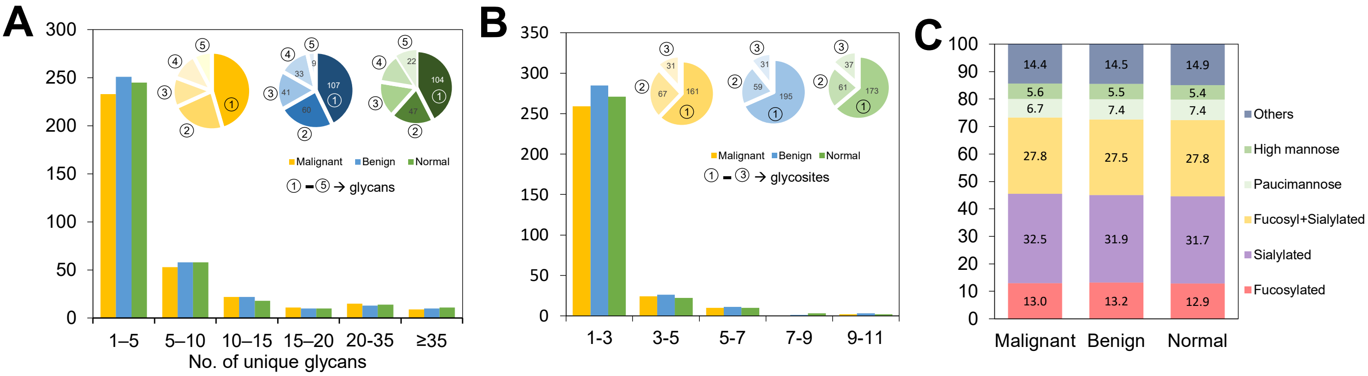


**Figure S3. (A)** Number of unique glycan per glycoprotein **(B)** Number of unique glycosites per glycoprotein **(C)** Glycan type distribution across clinical cohorts.
