## Supplementary material for "A Single-Aliquot, Enrichment-Free Workflow for High-Throughput Plasma Proteome and N-Glycoproteome Profiling": Figure S2

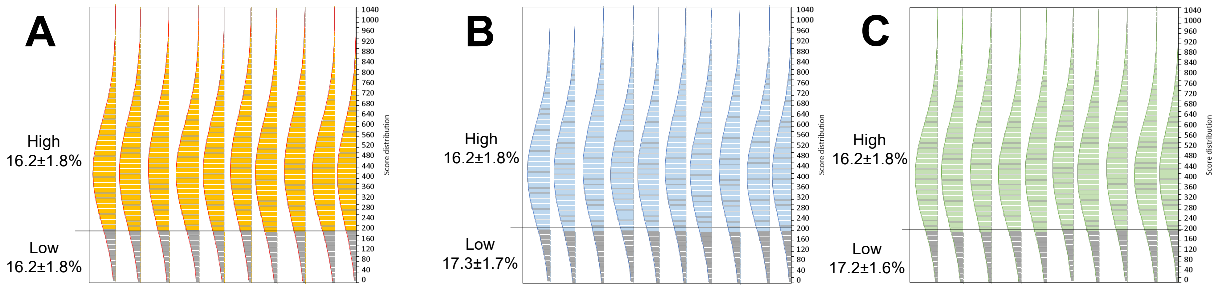


**Figure S2.** Byonic score distribution of unique glycopeptides in **(A)** malignant, **(B)** benign and **(C)** normal group.
