## Supplementary material for "A Single-Aliquot, Enrichment-Free Workflow for High-Throughput Plasma Proteome and N-Glycoproteome Profiling": Figure S1

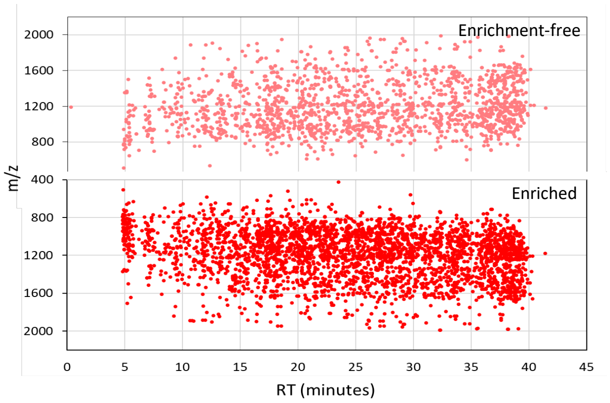


**Figure S1**. Comparison of glycopeptide distribution map between enrichment-free and enriched sample
