## Supplemental Table S5 for "A Single-Aliquot, Enrichment-Free Workflow for High-Throughput Plasma Proteome and N-Glycoproteome Profiling"

**Table S5.** List of significantly expressed glycopeptides across clinical groups (*cancer n=10, benign n=10, normal n=10*).

| No. | Peptide Sequence | Glycan | Protein | *p-value* |
|---|---|---|---|---|
| 1 | FSDGLESNSSTQFEVK | HexNAc(6)Hex(3) | C4A | 0.003501683 |
| 2 | ERSWPAVGNCSSALR | HexNAc(5)Hex(6)NeuAc(2) | HPX | 0.042326464 |
| 3 | VIDFNCTTSSVSSALANTK | HexNAc(4)Hex(5)Fuc(2)NeuAc(1) | HRG | 0.008827425 |
| 4 | TFVNITPAEVGVLVGK | HexNAc(5)Hex(3)Fuc(1) | PFN1 | 0.039247375 |
| 5 | PYLSAVRATLQAALCLENFSSQVVER | HexNAc(5)Hex(4) | ARPC4 | 0.020083396 |
| 6 | ALPQPQNVTSLLGCTH | HexNAc(4)Hex(5)Fuc(1)NeuAc(2) | HPX | 0.029354812 |
| 7 | LNAENNATFYFK | HexNAc(5)Hex(6)Fuc(3)NeuAc(2) | KNG1 | 0.039436292 |
| 8 | LNAENNATFYFK | HexNAc(4)Hex(5)Fuc(2)NeuAc(1) | KNG1 | 0.026613891 |
| 9 | NGSLFAFR | HexNAc(2)Hex(3) | VTN | 0.028585117 |
| 10 | ENLTAPGSDSAVFFEQGTTR | HexNAc(4)Hex(5)Fuc(1)NeuAc(1) | CP | 0.0264543 |
| 11 | TLNQSSDELQLSMGNAMFVK | HexNAc(5)Hex(6)Fuc(2) | SERPINA3 | 0.048514485 |
| 12 | CLQHFYGPNHEHCFNRTLLR | HexNAc(4)Hex(5)NeuAc(2) | MET | 0.036026345 |
| 13 | MLNTSSLLEQLNEQFNWVSR | HexNAc(4)Hex(6)Fuc(1)NeuAc(1) | CLU | 0.034546618 |
| 14 | TVLTPATNHMGNVTFTIPANREFK | HexNAc(2)Hex(6) | C3 | 0.043267612 |
| 15 | SIPACVPWSPYLFQPNDTCIVSGWGREK | HexNAc(2)Hex(5)Fuc(1) | CFI | 0.046085862 |
| 16 | FSDGLESNSSTQFEVK | HexNAc(6)Hex(4) | C4A | 0.02744594 |
| 17 | LSLHRPALEDLLLGSEANLTCTLTGLR | HexNAc(3)Hex(5)NeuAc(1) | IGHA1 | 0.045504618 |
| 18 | TVLTPATNHMGNVTFTIPANR | HexNAc(3)Hex(6) | C3 | 0.036628743 |
| 19 | SWPAVGNCSSALR | HexNAc(5)Hex(6)Fuc(2)NeuAc(1) | HPX | 0.033408201 |
| 20 | VQSTITSRMATTMIQSKVVNNSPQPQNVVFDVQIPK | HexNAc(2)Fuc(1) | ITIH2 | 0.044333866 |
| 21 | TKPREEQFNSTFR | HexNAc(5)Hex(4)Fuc(1) | IGHG2 | 0.038885529 |
| 22 | TTPKDFYVDENTTVR | HexNAc(5)Hex(4) | SERPINA4 | 0.038885529 |
| 23 | TKPREEQFNSTFR | HexNAc(4)Hex(5)Fuc(1)NeuAc(1) | IGHG2 | 0.009788929 |
| 24 | TKPREEQFNSTFR | HexNAc(4)Hex(5)Fuc(1) | IGHG2 | 0.004287122 |
| 25 | TTPKDFYVDENTTVR | HexNAc(4)Hex(4)NeuAc(1) | SERPINA4 | 0.006405636 |
| 26 | TKPREEQFNSTFR | HexNAc(4)Hex(4)Fuc(1)NeuAc(1) | IGHG2 | 0.006405636 |
| 27 | TKPREEQFNSTFR | HexNAc(4)Hex(4)Fuc(1) | IGHG2 | 0.005028784 |
| 28 | TTPKDFYVDENTTVR | HexNAc(4)Hex(4) | SERPINA4 | 0.03074102 |
| 29 | TKPREEQFNSTFR | HexNAc(3)Hex(3)Fuc(1) | IGHG2 | 0.009479975 |
| 30 | LHINHNNLTESVGPLPK | HexNAc(5)Hex(6)Fuc(1)NeuAc(1) | LUM | 0.037860574 |
| 31 | SLPNFPNTSATANATGGRALALQSSLPGAEGPDK | HexNAc(1)Fuc(1) | PI16 | 0.002304889 |
| 32 | VEIDTKSYWKALGISPFHEHAEVVFTANDSGPR | HexNAc(2)Hex(4)Fuc(1) | TTR | 0.031339595 |
| 33 | HEEGHMLNCTCFGQGR | HexNAc(4)Hex(5)Fuc(2)NeuAc(1) | FN1 | 0.037377656 |
| 34 | DQCIVDDITYNVNDTFHK | HexNAc(4)Hex(5)NeuAc(1) | FN1 | 0.016384445 |
| 35 | RHEEGHMLNCTCFGQGR | HexNAc(4)Hex(5) | FN1 | 0.024632577 |
| 36 | LNAENNATFYFK | HexNAc(5)Hex(6)NeuAc(3) | KNG1 | 0.043364137 |
| 37 | TAGWNVPIGTLRPFLNWTGPPEPIEAAVAR | HexNAc(2)Hex(1) | LTF | 0.009425632 |
| 38 | LNAENNATFYFK | HexNAc(4)Hex(5)NeuAc(1) | KNG1 | 0.011891763 |
| 39 | IYSGILNLSDITK | HexNAc(4)Hex(5)NeuAc(1) | KLKB1 | 0.00352347 |
| 40 | NFTENDLLVR | HexNAc(4)Hex(5)NeuAc(1) | F2 | 0.010439548 |
| 41 | LQAPLNYTEFQK | HexNAc(4)Hex(5)NeuAc(1) | KLKB1 | 0.027131731 |
| 42 | EVFVHPNYSK | HexNAc(4)Hex(5)NeuAc(2) | PROC | 0.014257911 |
| 43 | EYESYSDFERNVTEK | HexNAc(4)Hex(5)NeuAc(1) | C8B | 0.027215074 |
| 44 | SFEGLFYFLGSIVNFSQDPDVHFK | HexNAc(2) | CLTC | 0.02654147 |
| 45 | LVPHMNVSAVEK | HexNAc(5)Hex(6)NeuAc(3) | CNDP1 | 0.000924171 |
| 46 | PEINSTTHPGADLQENFCR | HexNAc(6)Hex(7)Fuc(1)NeuAc(1) | F2 | 0.032365694 |
| 47 | EVQQASLMFFVQLPSNTTWTLK | HexNAc(1)Fuc(1) | INHBC | 0.045997437 |
