## Supplemental Table S4 for "A Single-Aliquot, Enrichment-Free Workflow for High-Throughput Plasma Proteome and N-Glycoproteome Profiling"

**Table S4.** Orbitrap Astral Zoom mass spectrometer glycopeptide analysis parameters. (A) Global source and mass spectrometer parameters; (B) MS1 full scan experiment parameters; (C) ddMS2 scan experiment parameters.

**A**

| Global Parameters (Source & MS) | |
|---|---|
| Positive Ion Voltage | 2100 Volts |
| Ion Transfer Tube Temperature | 290°C |
| Expected Peak Width | 6 seconds |
| Default Charge State | 2 |
| Lock Mass Correction | Off |

**B**

| MS1 Full Scan Experiment Parameters | |
|---|---|
| Orbitrap Resolution | 180K |
| Scan Range (m/z) | 350-2000 |
| RF Lens (%) | 50 |
| Normalized AGC Target (%) / Absolute AGC Value | 500% / 5.00e6 |
| Maximum Injection Time | 50 milliseconds |
| Microscans | 1 |
| MIPS: Monoisotopic peak determination | Peptides |
| Charge State | 2-8 |
| Dynamic Exclusion Mode | Custom |
| Dynamic Exclusion Duration | 60 seconds |
| Dynamic Exclusion (after n times) | 1 |
| Dynamic Exclusion mass tolerance | (plus or minus) 10 ppm |
| Dynamic Exclusion isotope exclusion | yes |

**C**

| ddMS2 Scan Experiment Parameters | |
|---|---|
| Scan Mass Range (m/z) | 120-2000 |
| Isolation Window (m/z) | 2 |
| Isolation offset | Off |
| AGC Target | Custom |
| Normalized AGC Target (%) / Absolute AGC Value | 100% / 1.00e4 |
| Maximum Injection Time | 10 milliseconds |
| HCD Collision Energy (%) | 24,32,40 |
| RF Lens (%) | 40 |
| Detector | Astral |
| Microscans | 1 |
| Data Dependent Mode | Cycle Time |
| Data Dependent Mode: Time in between master scans | 0.5 seconds |
